# Anthropogenic risk factors as the major cause of cancer driver events

**DOI:** 10.1101/486860

**Authors:** Aleksey V. Belikov

## Abstract

**Background:** I have recently shown that the number of rate-limiting driver events per tumor can be estimated from the age distribution of cancer incidence using the gamma/Erlang probability distribution. It is important to understand how these predictions relate to established risk factors.

**Methods:** The number of rate-limiting driver events per tumor was estimated using the gamma/Erlang distribution and correlated to the percentage of cancer cases attributable to modifiable risk factors.

**Results:** The predicted number of rate-limiting driver events per tumor strongly correlates with the proportion of cancer cases attributable to modifiable risk factors for all cancers except those induced by infection or ultraviolet radiation. The correlation was confirmed for three countries, three corresponding incidence databases and risk estimation studies, as well as for both sexes: USA, males [r=0.80, P=0.002], females [r=0.81, P=0.0003]; England, males [r=0.90, P<0.0001], females [r=0.67, P=0.002]; Australia, males [r=0.90, P=0.0004], females [r=0.68, P=0.01].

**Conclusions:** It is thus confirmed that predictions based on interpreting the age distribution of cancer incidence as the gamma/Erlang probability distribution have biological meaning, validating the underlying Poisson process as the law governing the development of the majority of cancer types, especially those driven by chemical mutagens. Importantly, this study suggests that the majority of driver events (60-80% in males, 50-70% in females) are induced by anthropogenic carcinogens, and not by cell replication errors or other internal processes.

## Introduction

There have been multiple attempts to deduce the number of rate-limiting steps in carcinogenesis from the age distribution of cancer incidence or mortality [1]. The proposed models for doing this, however, suffer from several serious drawbacks. For example, early models assumed that cancer mortality increases with age according to the power law [2-4], which is inconsistent with the observed deceleration of mortality growth at an advanced age. Moreover, when high quality data have accumulated, it became clear that, at least for some cancers, incidence even starts to decrease after peaking at some advanced age [5, 6]. More recent models of cancer progression are based on multiple biological assumptions, consist of complicated equations that incorporate many predetermined empirical parameters, and still have not been shown to describe the decrease in cancer incidence at an advanced age [7-12]. It is also clear that an infinite number of such mechanistic models can be created and custom tailored to fit any set of data, leading us to question their explanatory and predictive values.

I have recently proposed that the age distribution of cancer incidence can be interpreted as the statistical distribution of probability to accumulate the required number of driver events by the given age [13]. I have shown that, of all standard probability distributions, the gamma distribution (and its special case with the integer shape parameter – the Erlang distribution) fits the actual age distribution of incidence for 20 most prevalent cancers the best [13]. I have then shown that the gamma/Erlang distribution is the only standard distribution that, in addition, approximates incidence for all studied childhood and young adulthood cancers, thus validating it as the universal equation describing cancer incidence [14]. Importantly, the Erlang distribution describes the waiting time for the occurrence of the given number of independent random events, as it was initially devised to calculate call queues at telephone exchanges. It is based on the Poisson process, which implies not only pure randomness of event timings but also their constant average rate. Thus, the excellent fit of the gamma/Erlang distribution to the actual incidence data implies that cancers develop according to the Poisson process, i.e. driver events occur randomly and at a constant average rate.

Interestingly, the shape parameter of the gamma/Erlang distribution can be interpreted as the number of rate-limiting driver events that occur by the time of cancer diagnosis. It is thus possible to estimate this number for any cancer type, upon fitting the gamma/Erlang distribution to the actual age distribution of incidence. I have shown that these numbers vary considerably, from 1 in retinoblastoma [14] to 41 in prostate cancer [13]. Next, it is important to show that these predictions correspond to experimentally observed variables, such as the number of driver mutations per tumor predicted from sequencing data. However, the variability of DNA alterations that can contribute to cancer progression, some of which are not yet routinely assessed, and the imperfection of algorithms for separating driver and passenger mutations severely complicate this task, as discussed in [13]. Thus, a simpler correlate is required to prove the meaningfulness of the predictions, before engaging in a full-scale confirmation effort.

Here I identify such correlate as the percentage of cancer cases due to modifiable risk factors. This is an often-used parameter in epidemiological studies, and is also called the population attributable fraction (PAF). It shows, for example, what percentage of lung cancer cases are caused by smoking tobacco. Combined PAF shows the overall contribution of all potentially modifiable risk factors, which usually include air pollution, occupational hazards, ionizing radiation, smoking, alcohol, poor diet, insufficient exercise, obesity, infection and ultraviolet radiation. Here I show that the numbers of driver events per tumor predicted by the gamma/Erlang distribution strongly correlate with combined PAFs for most cancers, with the exception of cancers with the large contribution from infection or ultraviolet radiation. This confirms that predictions obtained from the gamma/Erlang distribution are meaningful, validating the Poisson process as the law governing the development of most cancer types and fostering the search for correlations with tumor sequencing data. Importantly, the results suggest that up to 80% of driver events are caused by the environment and lifestyle, and not, for example, by stem cell divisions, as has been recently proposed [15, 16].

## Methods

### I. Data acquisition

#### a) Population attributable fractions data

Population attributable fractions (PAFs) combining all risk factors were obtained directly from published open-access articles separately for each cancer type and sex. PAFs for USA were obtained from the publication by Islami et al., Table 2 (Ref[17]). PAFs for England were obtained from the publication by Brown et al., Table 2 (Ref[18]). PAFs for Australia were obtained from the publication by Whiteman et al., Table 2 (Ref[19]). No modification or processing of PAF data was performed.

**Table 1.** Goodness of fit (R^2^) of the gamma distribution to the actual cancer incidence data

| Cancer type | England |  | USA |  | Australia |  |
| --- | --- | --- | --- | --- | --- | --- |
|  | Males | Females | Males | Females | Males | Females |
| Lung | 0.9996 | 0.9989 | 0.9990 | 0.9973 | 0.9997 | 0.9994 |
| Uterus | - | 0.9965 | - | 0.9954 | - | 0.9934 |
| Mesothelioma | 0.9997 | 0.9980 | ND | ND | ND | ND |
| Larynx | 0.9989 | 0.9964 | 0.9997 | 0.9945 | 0.9965 | 0.9838 |
| Pancreas | 0.9999 | 0.9999 | 1.000 | 0.9998 | 0.9995 | 0.9982 |
| Bladder | 0.9998 | 0.9998 | 0.9997 | 0.9996 | 0.9998 | 0.9994 |
| Myeloma | 0.9999 | 0.9998 | 0.9994 | 0.9992 | ND | ND |
| Kidney | 0.9991 | 0.9986 | 0.9985 | 0.9959 | 0.9970 | 0.9985 |
| Gallbladder | 0.9998 | 0.9991 | 0.9994 | 0.9996 | 0.9996 | 0.9992 |
| Ovary | - | 0.9990 | - | 0.9990 | - | 0.9956 |
| Colorectum | 0.9998 | 0.9997 | 0.9993 | 0.9987 | 0.9987 | 0.9987 |
| Liver* | 0.9987 | 0.9994 | 0.9839 | 0.9985 | 0.9797 | 0.9975 |
| Esophagus | 0.9997 | 0.9997 | 0.9999 | 0.9991 | 0.9989 | 0.9981 |
| Non-Hodgkin lymphoma | 0.9970 | 0.9976 | 0.9973 | 0.9969 | 0.9985 | 0.9974 |
| Oral cavity | 0.9912 | 0.9973 | ND | ND | ND | ND |
| Oral cavity and pharynx | ND | ND | 0.9971 | 0.9987 | 0.9972 | 0.9966 |
| Breast | - | 0.9828 | - | 0.9978 | - | 0.9887 |
| Brain | 0.9797 | 0.9775 | ND | ND | ND | ND |
| Leukemia | 0.9957 | 0.9961 | ND | ND | 0.9950 | 0.9912 |
| Myeloid leukemia | ND | ND | 0.9930 | 0.9891 | ND | ND |
| Thyroid | 0.9859 | 0.9251 | 0.9808 | 0.9870 | ND | ND |
| Melanoma | 0.9970 | 0.9962 | 0.9987 | 0.9945 | 0.9983 | 0.9984 |
| Stomach | 0.9988 | 0.9983 | 0.9993 | 0.9979 | 0.9999 | 0.9965 |
| Anus* | 0.9982 | 0.9934 | 0.9927 | 0.9827 | 0.9926 | 0.9700 |
| Pharynx* | 0.9848 | 0.9790 | ND | ND | ND | ND |
| Nasopharynx* | 0.9776 | 0.8836 | ND | ND | ND | ND |
| Kaposi sarcoma* | 0.6761 | 0.3861 | 0.1997 | 0.9906 | 0.7562 | 0.9942 |
| Hodgkin lymphoma* | 0.5301 | 0.1664 | 0.5702 | 0.1730 | 0.4513 | 0.1200 |
| Penis* | 0.9961 | - | 0.9982 | - | 0.9762 | - |
| Cervix* | - | 0.6003 | - | 0.9033 | - | 0.8016 |
| Vagina* | - | ND | - | 0.9985 | - | 0.9918 |
| Vulva* | - | ND | - | 0.9900 | - | 0.9773 |

ND – no incidence data in the database or no corresponding PAF data in the source publication.
Asterisk (\*) denotes cancers in which viral infection contributes to more than 30% of cases, according to the published PAF data [17-19].

**Table 2.** Predicted numbers of driver events per tumor and estimated percentages of cases due to anthropogenic risk factors for cancers in males.

| Cancer type | England |  | USA |  | Australia |  |
| --- | --- | --- | --- | --- | --- | --- |
|  | Predicted number of driver events per tumor | Estimated percentage of cases due to modifiable risk factors [17] | Predicted number of driver events per tumor | Estimated percentage of cases due to modifiable risk factors [18] | Predicted number of driver events per tumor | Estimated percentage of cases due to modifiable risk factors [19] |
| Mesothelioma | 29 | 97 | - | ND | - | ND |
| Lung | 24 | 82 | 28 | 89 | 22 | 86 |
| Larynx | 20 | 73 | 22 | 84 | 21 | 84 |
| Bladder | 20 | 51 | 20 | 49 | 14 | 34 |
| Colorectum | 18 | 57 | 12 | 58 | 18 | 56 |
| Oral cavity | 16 | 53 | - | ND | - | ND |
| Liver | 15 | 53 | - | NA | - | NA |
| Esophagus | 14 | 61 | 19 | 75 | 14 | 74 |
| Pancreas | 14 | 34 | 15 | 26 | 12 | 31 |
| Kidney | 13 | 32 | 15 | 52 | 12 | 39 |
| Myeloma | 12 | 16 | 16 | 11 | - | ND |
| Gallbladder | 12 | 13 | 18 | 33 | 11 | 16 |
| Brain | 9 | 0 | - | ND | - | ND |
| Leukemia | 9 | 11 | - | ND | 8 | 7 |
| Myeloid leukemia | - | ND | 8 | 17 | - | ND |
| Non-Hodgkin lymphoma | 8 | 3 | 7 | 14 | 7 | 4 |
| Thyroid | 3 | 10 | 6 | 12 | - | ND |

ND – no data in the source publication, NA – assigned to the non-anthropogenic group due to the strong contribution of the viral infection.

#### b) USA incidence data

United States Cancer Statistics Public Information Data: Incidence 1999–2012 was downloaded from the Centers for Disease Control and Prevention Wide-ranging OnLine Data for Epidemiologic Research (CDC WONDER) online database (http://wonder.cdc.gov/cancer-v2012.HTML) in November 2018 (Ref[20]). The United States Cancer Statistics (USCS) are the official federal statistics on cancer incidence from registries having high-quality data for 50 states and the District of Columbia. Data are provided by The Centers for Disease Control and Prevention National Program of Cancer Registries (NPCR) and The National Cancer Institute Surveillance, Epidemiology and End Results (SEER) program. Results were grouped by 5-year Age Groups and Crude Rates were selected as output. Crude Rates are calculated as the number of new cancer cases reported each calendar year per 100,000 population in each 5-year age group. The data were downloaded separately for males and females for each cancer type listed in the publication by Islami et al., Table 2 (Ref[17]).

#### c) England incidence data

England cancer incidence data were downloaded from the European Cancer Information System (ECIS) Data explorer (https://ecis.jrc.ec.europa.eu/explorer.php?$0-1$1-UK$2-224$4-1,2$3-All$6-5,84$5-1999,2012$7-2$CRatesByCancer$X0_10-ASR_EU_NEW) in November 2018 (Ref[21]). The ECIS database contains the aggregated output and the results computed from data submitted by population-based European cancer registries participating in Europe to the European Network of Cancer Registries – Joint Research Centre (ENCR-JRC) project on “Cancer Incidence and Mortality in Europe". Years of observation were limited to 1999-2012 period, to match the USA data. Incidence is calculated as the number of new cancer cases reported each calendar year per 100,000 population in each 5-year age group. The data were downloaded separately for males and females for each cancer type listed in the publication by Brown et al., Table 2 (Ref[18]), except for vulva and vagina cancers, as their selection was not possible in ECIS Data explorer.

#### d) Australia incidence data

Australia cancer incidence data were downloaded from the Cancer Incidence in Five Continents (CI5) Volume XI Age-specific curves Online Analysis tool (http://ci5.iarc.fr/CI5-XI/Pages/age-specific-curves_sel.aspx) in November 2018 (Ref[22]). CI5 is published approximately every five years by the International Agency for Research on Cancer (IARC) and the International Association of Cancer Registries (IACR) and provides comparable high quality statistics on the incidence of cancer from cancer registries around the world. Volume XI contains information from 343 cancer registries in 65 countries for cancers diagnosed from 2008 to 2012. Incidence is calculated as the number of new cancer cases reported each calendar year per 100,000 population in each 5-year age group. The data were downloaded separately for males and females for each cancer type listed in the publication by Whiteman et al., Table 2 (Ref[19]).

### II. Data selection and analysis

#### a) Estimation of the number of driver events per tumor

For analysis, the incidence data were imported into GraphPad Prism 6. The following age groups were selected: “5–9 years”, “10–14 years”, “15–19 years”, “20–24 years”, “25–29 years”, “30–34 years”, “35–39 years”, “40–44 years”, “45– 49 years”, “50–54 years”, “55–59 years”, “60–64 years “, “65–69 years”, “70–74 years”, “75–79 years” and “80–84 years”. Prior age groups were excluded due to possible contamination by childhood subtype incidence, and “85+ years” was excluded due to an undefined age interval. If in the first several age groups (“5–9 years”, “10–14 years”, “15–19 years”) incidence initially decreased with age, reflecting contamination by childhood subtype incidence, these values were removed until a steady increase in incidence was detected. The middle age of each age group was used for the x values, e.g. 17.5 for the “15–19 years” age group. Incidence (new cancer cases per calendar year per 100,000 population) for each age group and each cancer type was used for the y values. Data for different countries, as well as for males and females, were analyzed separately. Data were analyzed with Nonlinear regression using the following User-defined equation for the gamma distribution:

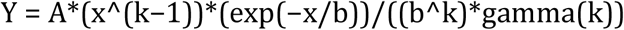

The amplitude parameter *A* was constrained to “Must be between zero and 100000.0” and scale and shape parameters *b* and *k* to “Must be greater than 0.0”. “Initial values, to be fit” for all parameters were set to 1.0. All other settings were kept at default values, e.g. Least squares fit and No weighting.

The numerical value of the shape parameter *k* rounded to the nearest integer is interpreted as the number of driver events per tumor [13].

#### b) Correlation of the predicted numbers of driver events per tumor with PAFs

Obtained *k* values were correlated to population attributable fractions (PAFs) in GraphPad Prism 6 using the inbuilt Correlation tool at default settings, e.g. Pearson correlation with two-tailed P value. Cancer types were sorted into two classes, and correlation was performed separately for each class. Cancer types in which infection (*Helicobacter pylori*, Hepatitis B virus, Hepatitis C virus, Human herpes virus type 8: Kaposi sarcoma herpes virus, Human immunodeficiency virus and Human papillomavirus) or ultraviolet radiation contributed to more than 30% of cases, for a given country according to the published PAF data [17-19], were assigned to Class 2 (non-anthropogenic). The rest were assigned to Class 1 (anthropogenic), which included cancers with substantial contribution from air pollution, occupational exposure, exposure to ionizing radiation, smoking and exposure to secondhand smoke, alcohol intake, poor diet (red and processed meat, insufficient fiber, vegetables, fruit and calcium), excess body weight, insufficient physical activity, insufficient breastfeeding, postmenopausal hormone therapy and oral contraceptives, according to the published PAF data [17-19].

## Results

To estimate the numbers of driver events per tumor, the gamma distribution was fitted to the actual age distributions of incidence separately for males and females in three countries – USA, England and Australia (Figure 1 and Table 1). The fits were generally excellent (R^2^=0.99), except for brain cancer (R^2^=0.98), thyroid cancer (R^2^=0.97), and several virus-induced cancers: pharyngeal (R^2^=0.98), nasopharyngeal (R^2^=0.93), vulvar (R^2^=0.98), cervical (R^2^=0.77), Kaposi sarcoma (R^2^=0.67) and Hodgkin lymphoma (R^2^=0.34). Due to the unsatisfactory fits, the last three cancer types were excluded from the further analysis. Successful fitting of the remaining cancer types allowed the estimation of the numbers of driver events per tumor using the shape parameter of the gamma distribution.

**Figure 1.**
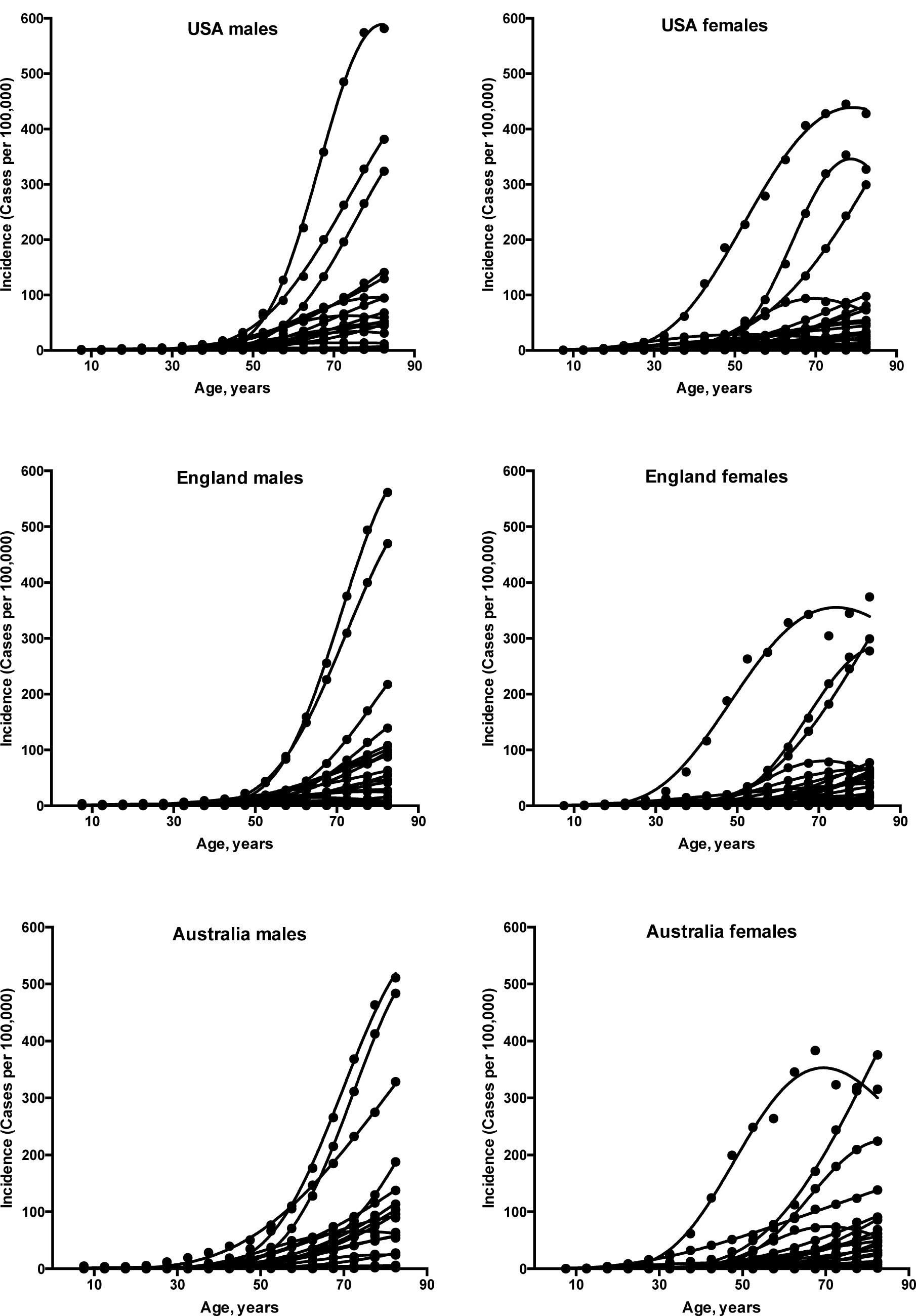
Fits of the gamma distribution to the actual incidence data for various cancers

Plotting the correlation of the number of driver events per tumor predicted from the gamma distribution with the estimated percentage of cases due to modifiable risk factors obtained from the published studies revealed that cancers appear to cluster in two classes. Class 1, which included the majority of cancers, demonstrated the linear correlation, whereas Class 2 clustered in the upper left corner of the plot in a cloud-like fashion. Investigation of the Class 2 revealed that it consists entirely of cancers with substantial (>30%) contribution of infection to their pathogenesis, plus the melanoma cancer. Class 2 was therefore named “non-anthropogenic”, as infections and ultraviolet radiation existed long before the human civilization. Interestingly, all cancers in Class 1 were induced by factors that arose with human civilization, such as air pollution, occupational hazards, ionizing radiation, smoking, alcohol, poor diet, insufficient exercise, obesity, insufficient breastfeeding, postmenopausal hormone therapy and oral contraceptives. Therefore, Class 1 was termed “anthropogenic”.

The correlation of the predicted number of driver events per tumor with the estimated percentage of cases due to modifiable risk factors for cancers in males is shown in Figure 2 and Table 2, and in females in Figure 3 and Table 3. It can be seen that anthropogenic cancers indeed exhibit the strong correlation for all studied countries and for both sexes, whereas non-anthropogenic cancers exhibit the correlation in none of the cases. Amongst anthropogenic cancers, the correlation is stronger and more significant for males than for females. Interestingly, the correlation is stronger and more significant for American females [r=0.81, P=0.0003] than for English [r=0.67, P=0.002] and Australian [r=0.68, P=0.01] females, but weaker and less significant for USA males [r=0.80, P=0.002] than for English [r=0.90, P<0.0001] and Australian [r=0.90, P=0.0004] males. These differences are likely explained by differing exposures to risk factors between countries and between sexes, as well as by variations in the screening, diagnostics and reporting protocols of different countries, in the sets of cancers included in the studies from which risk factor data were obtained, and in the methodologies of those studies. The role of population genetics also cannot be ruled out.

**Table 3.** Predicted numbers of driver events per tumor and estimated percentages of cases due to anthropogenic risk factors for cancers in females.

| Cancer type | England |  | USA |  | Australia |  |
| --- | --- | --- | --- | --- | --- | --- |
|  | Predicted number of driver events per tumor | Estimated percentage of cases due to modifiable risk factors [17] | Predicted number of driver events per tumor | Estimated percentage of cases due to modifiable risk factors [18] | Predicted number of driver events per tumor | Estimated percentage of cases due to modifiable risk factors [19] |
| Lung | 23 | 75 | 30 | 83 | 22 | 79 |
| Uterus | 23 | 34 | 20 | 71 | 22 | 33 |
| Mesothelioma | 22 | 83 | - | ND | - | ND |
| Larynx | 18 | 66 | 25 | 79 | 28 | 78 |
| Pancreas | 15 | 29 | 15 | 25 | 15 | 28 |
| Bladder | 14 | 43 | 17 | 39 | 11 | 26 |
| Myeloma | 14 | 11 | 16 | 12 | - | ND |
| Kidney | 13 | 36 | 14 | 56 | 10 | 24 |
| Gallbladder | 12 | 23 | 14 | 37 | 15 | 13 |
| Ovary | 13 | 11 | 8 | 4 | 7 | 7 |
| Colorectum | 12 | 51 | 9 | 51 | 11 | 42 |
| Liver | 12 | 39 | - | NA | - | NA |
| Esophagus | 11 | 54 | 17 | 68 | 11 | 76 |
| Non-Hodgkin lymphoma | 11 | 3 | 10 | 2 | 7 | 3 |
| Oral cavity | 9 | 34 | - | ND | - | ND |
| Breast | 9 | 23 | 10 | 29 | 11 | 23 |
| Brain | 8 | 5 | - | ND | - | ND |
| Leukemia | 7 | 13 | - | ND | 8 | 2 |
| Myeloid leukemia | - | ND | 6 | 13 | - | ND |
| Thyroid | 3 | 9 | 5 | 13 | - | ND |

ND – no data in the source publication, NA – assigned to the non-anthropogenic group due to the strong contribution of the viral infection.

**Figure 2.**
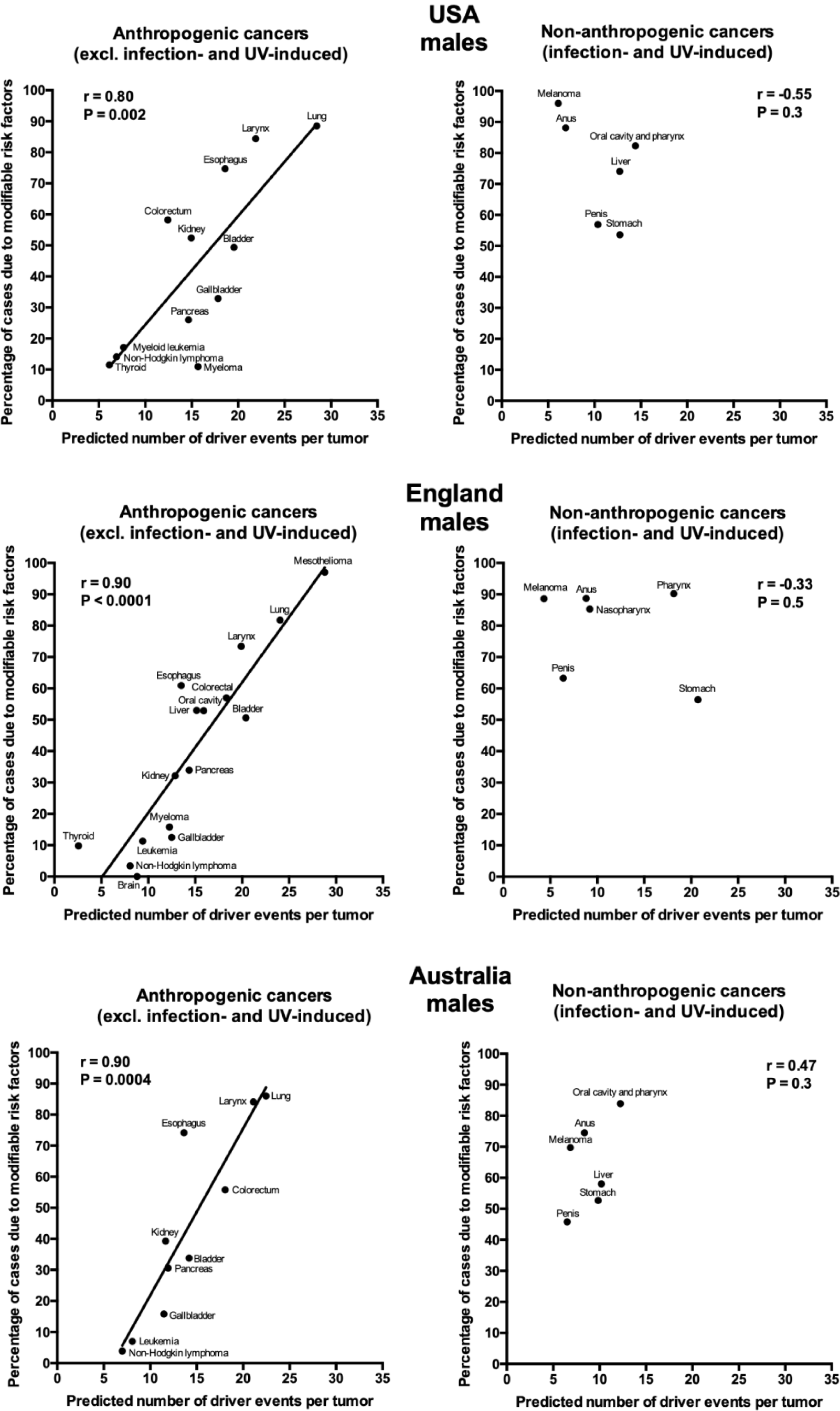
Correlation of the predicted numbers of driver events per tumor with the estimated percentages of cases due to modifiable risk factors for cancers in males.

**Figure 3.**
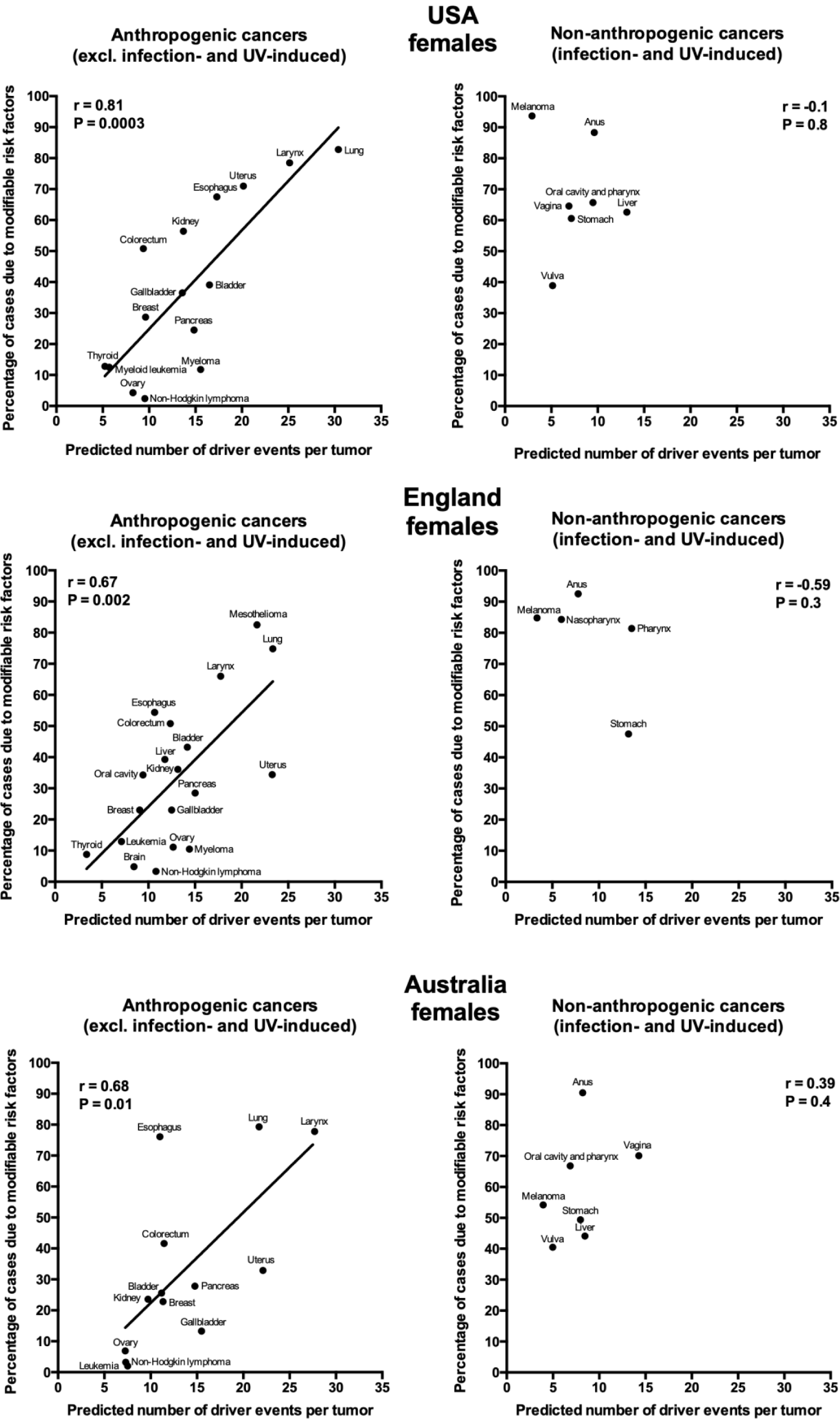
Correlation of the predicted numbers of driver events per tumor with the estimated percentages of cases due to modifiable risk factors for cancers in females.

## Discussion

One of the most interesting findings of this study is the clustering of all cancers into two classes, termed here anthropogenic and non-anthropogenic. The possible explanation for this dichotomy is that *the human body managed to evolve some protective countermeasures against cancer risk factors that were present for millions of years, whereas it appears unprepared for the novel risk factors brought by our civilization*. For example, ultraviolet radiation has been present on Earth since the beginning, and although melanocytes cannot completely protect their DNA, and a lot of DNA damage occurs, it is likely that they developed a very slow division rate [23] to avoid conversion of this damage into mutations for as long as possible. This may explain why only few rate-limiting driver events are predicted for melanoma despite lots of DNA damage that melanocytes receive – rate-limiting in this case is cell division and not the DNA damage. Similarly, the human body had plenty of time to adapt to viruses and install some blocks which are difficult for viruses to overcome, which may explain why the incidence rates of virus-induced cancers are low, and less driver events are predicted than would be expected from the linear correlation. It is also clear that viruses are inducing cancer via different mechanisms than chemical carcinogens [24, 25], and thus the development of such cancers may not be described by the Poisson process. Indeed, many of the virus-induced cancers have rather poor fits of the Erlang distribution to their age distributions of incidence (Table 1).

The strong positive correlation of the predicted number of driver events per tumor with the contribution from anthropogenic risk factors suggests that the majority of driver events are caused by those factors. In other words, *the higher is the number of driver events that are required for a given cancer type to appear, the less likely is for them to occur by chance (e.g. due to replication errors), and the more dependent are they on anthropogenic carcinogens to be induced*. Indeed, as r^2^ is called “the coefficient of determination” and describes the proportion of the variance in one variable that is explained by the other variable, we can calculate (by squaring Pearson r values from Figures 2 and 3) that anthropogenic risk factors explain 64%, 81% and 81% of the variance in the predicted number of driver events per tumor for males and 66%, 45% and 46% of the variance for females, living in USA, England and Australia, respectively. This is in accord with the mainstream view that the environment and lifestyle are the major contributors to carcinogenesis, but conflicts with the recently proposed view that the majority of cancers develop due to replicative mutations occurring during stem cell division [15, 16]. The latter view is based on predominantly mouse data handpicked from varied publications and processed through calculations with unobvious assumptions, and thus has been widely criticized [26-32].

It is also interesting to speculate why the observed correlations are stronger for males than for females. One likely explanation is that males generally are more exposed to chemical mutagens, e.g. during smoking and at dangerous industries [17-19], directly inducing mutations in the DNA, some of which happen to be drivers. On the other hand, females have a higher contribution to cancer risk from disturbances in physiology, usually related to hormone levels, such as being obese, using oral contraceptives, undergoing postmenopausal hormone therapy or abstaining from breastfeeding [17-19]. *These risk factors may not lead to an increase in the number of overall and driver mutations (discrete events), but promote cancer via changes in intracellular signaling levels or the microenvironment (gradual change)* [33-37]. The latter cannot be detected and counted using the gamma/Erlang distribution, which is capable of recognizing only discrete random events.

Overall, the correlations identified here serve as the validation of the hypothesis that most cancers develop according to the Poisson process and that the gamma/Erlang distribution can be used to predict the number of driver events per tumor for most cancer types, especially those driven by chemical mutagens [13, 14]. This has numerous implications, from the fundamental understanding of the carcinogenesis process to the improvement in driver prediction algorithms.

## References

1. Hornsby C, Page KM, Tomlinson IP. What can we learn from the population incidence of cancer? Armitage and Doll revisited. Lancet Oncol 2007;8(11):1030–8.

2. Nordling CO. A new theory on cancer-inducing mechanism. Br J Cancer 1953;7(1):68–72.

3. Armitage P, Doll R. The age distribution of cancer and a multi-stage theory of carcinogenesis. Br J Cancer 2004;91(12):1983–9.

4. Knudson AG. Two genetic hits (more or less) to cancer. Nat Rev Cancer 2001;1(2):157–62.

5. Saltzstein SL, Behling CA, Baergen RN. Features of cancer in nonagenarians and centenarians. J Am Geriatr Soc 1998;46(8):994–8.

6. Harding C, Pompei F, Wilson R. Peak and decline in cancer incidence, mortality, and prevalence at old ages. Cancer 2012;118(5):1371–86.

7. Luebeck EG, Moolgavkar SH. Multistage carcinogenesis and the incidence of colorectal cancer. Proc Natl Acad Sci U S A 2002;99(23):15095–100.

8. Little MP, Wright EG. A stochastic carcinogenesis model incorporating genomic instability fitted to colon cancer data. Math Biosci 2003;183(2):111–34.

9. Michor F, Iwasa Y, Nowak MA. The age incidence of chronic myeloid leukemia can be explained by a one-mutation model. Proc Natl Acad Sci U S A 2006;103(40):14931–4.

10. Meza R, Jeon J, Moolgavkar SH, et al. Age-specific incidence of cancer: Phases, transitions, and biological implications. Proc Natl Acad Sci U S A 2008;105(42):16284–9.

11. Calabrese P, Shibata D. A simple algebraic cancer equation: calculating how cancers may arise with normal mutation rates. BMC Cancer 2010;10:3.

12. Luebeck EG, Curtius K, Jeon J, et al. Impact of tumor progression on cancer incidence curves. Cancer Res 2013;73(3):1086–96.

13. Belikov AV. The number of key carcinogenic events can be predicted from cancer incidence. Sci Rep 2017;7(1):12170.

14. Belikov AV. The Poisson process is the universal law of cancer development: driver mutations accumulate randomly, silently, at constant rate and for many decades, likely in stem cells. bioRxiv, 2018; https://doi.org/10.1101/231027, (posted December 04, 2018).

15. Tomasetti C, Vogelstein B. Cancer etiology. Variation in cancer risk among tissues can be explained by the number of stem cell divisions. Science 2015;347(6217):78–81.

16. Tomasetti C, Li L, Vogelstein B. Stem cell divisions, somatic mutations, cancer etiology, and cancer prevention. Science 2017;355(6331):1330–1334.

17. Islami F, Goding Sauer A, Miller KD, et al. Proportion and number of cancer cases and deaths attributable to potentially modifiable risk factors in the United States. CA Cancer J Clin 2018;68(1):31–54.

18. Brown KF, Rumgay H, Dunlop C, et al. The fraction of cancer attributable to modifiable risk factors in England, Wales, Scotland, Northern Ireland, and the United Kingdom in 2015. Br J Cancer 2018;118(8):1130–1141.

19. Whiteman DC, Webb PM, Green AC, et al. Cancers in Australia in 2010 attributable to modifiable factors: summary and conclusions. Aust N Z J Public Health 2015;39(5):477–84.

20. United States Cancer Statistics: 1999 - 2012 Archive Incidence, WONDER Online Database. http://wonder.cdc.gov/cancer-v2012.html United States Department of Health and Human Services, Centers for Disease Control and Prevention and National Cancer Institute; 2015.

21. ECIS - European Cancer Information System. https://ecis.jrc.ec.europa.eu European Union; 2018.

22. Bray F CM, Mery L, Piñeros M, Znaor A, Zanetti R and Ferlay J, editors. Cancer Incidence in Five Continents, Vol. XI (electronic version). http://ci5.iarc.fr International Agency for Research on Cancer; 2017.

23. Halaban R, Ghosh S, Duray P, et al. Human melanocytes cultured from nevi and melanomas. J Invest Dermatol 1986;87(1):95–101.

24. Butel JS. Viral carcinogenesis: revelation of molecular mechanisms and etiology of human disease. Carcinogenesis 2000;21(3):405–26.

25. Mesri EA, Feitelson MA, Munger K. Human viral oncogenesis: a cancer hallmarks analysis. Cell Host Microbe 2014;15(3):266–82.

26. Giovannucci EL. Are Most Cancers Caused by Specific Risk Factors Acting on Tissues With High Underlying Stem Cell Divisions? J Natl Cancer Inst 2015;108(3).

27. Rozhok AI, Wahl GM, DeGregori J. A Critical Examination of the “Bad Luck” Explanation of Cancer Risk. Cancer Prev Res (Phila) 2015;8(9):762–4.

28. Wu S, Powers S, Zhu W, et al. Substantial contribution of extrinsic risk factors to cancer development. Nature 2016;529(7584):43–7.

29. Wensink MJ, Vaupel JW, Christensen K. Stem Cell Divisions Per Se Do Not Cause Cancer. Epidemiology 2017;28(4):e35–e37.

30. Ashford NA, Bauman P, Brown HS, et al. Cancer risk: role of environment. Science 2015;347(6223):727.

31. Gotay C, Dummer T, Spinelli J. Cancer risk: prevention is crucial. Science 2015;347(6223):728.

32. O’Callaghan M. Cancer risk: accuracy of literature. Science 2015;347(6223):729.

33. Tahergorabi Z, Khazaei M, Moodi M, et al. From obesity to cancer: a review on proposed mechanisms. Cell Biochem Funct 2016;34(8):533–545.

34. Quail DF, Dannenberg AJ. The obese adipose tissue microenvironment in cancer development and progression. Nat Rev Endocrinol 2019;15(3):139–154.

35. McNamara KM, Guestini F, Sakurai M, et al. How far have we come in terms of estrogens in breast cancer? [Review]. Endocr J 2016;63(5):413–24.

36. Carroll JS. Mechanisms of oestrogen receptor (ER) gene regulation in breast cancer. Eur J Endocrinol 2016;175(1):R41–9.

37. Trevino LS, Wang Q, Walker CL. Hypothesis: Activation of rapid signaling by environmental estrogens and epigenetic reprogramming in breast cancer. Reprod Toxicol 2015;54:136–40.

